# ToxiTaRGET: a multi-omics resource for toxicant-responsive molecular targets

**DOI:** 10.1101/2025.07.28.667228

**Authors:** Ravindra Kumar, Tianyi Fu, Prashant Kumar Kuntala, Benpeng Miao, Shuhua Fu, Daofeng Li, Marisa S. Bartolomei, Cheryl Walker, Ting Wang, Bo A. Zhang

## Abstract

Environmental toxicant exposures can induce widespread alterations in both the transcriptome and epigenome of mammals, and directly contribute to the increased risk of various diseases, including cardiovascular disorders, cancer, and neurological disorders. To evaluate how early-life toxicants produce long-term impacts on the transcriptome and epigenome in mice, the Toxicant Exposures and Responses by Genomic and Epigenomic Regulators of Transcription II (TaRGET II) Consortium generated a landmark resource comprising 3,607 multi-omics from longitudinal studies in mice. The molecular changes in responding to distinct environmental toxicants, including arsenic (As), lead (Pb), bisphenol A (BPA), tributyltin (TBT), di-2-ethylhexyl phthalate (DEHP), dioxin (TCDD), and fine particulate matter (PM2.5), were systematically identified and visualized on an integrative platform, ToxiTaRGET, to allow quickly search and browse by researchers. ToxiTaRGET houses a rich repository of molecular signatures, including gene expression, chromatin accessibility, and DNA methylation profiles, in response to early-life toxicant exposures. These molecular signatures span multiple biologically important tissues in both male and female mice at three distinct life stages, offering a valuable resource for the environmental health and toxicogenomic research communities.

## Introduction

In recent decades, the burden of diseases linked to environmental exposures has increased significantly^1-3^. Among various environmental hazards, toxicants are particularly dangerous, as they can enter the body through inhalation, ingestion, or dermal absorption^4-7^. Once inside, these chemicals interact with critical biomolecules such as DNA, RNA, proteins, lipids, and metabolites, leading to structural and functional disruptions^8,9^. These molecular alterations can impair cellular homeostasis and are implicated in the development of numerous chronic diseases, including cancer, metabolic syndrome, cardiovascular disease, and neurological disorders^9-14^. To better understand how specific environmental toxicants affect human health and influence molecular targets, distinct technologies, particularly recently developed omics approaches, enable high-throughput profiling of molecular changes in response to toxicant exposure^15-19^. High-throughput technologies, such as transcriptomics, epigenomics, proteomics, and metabolomics, have been widely used and provide unprecedented opportunities to understand how toxicants disturb cellular pathways and lead to disease^20-22^.

To comprehensively uncover the molecular mechanisms driving transcriptomic and epigenomic responses to environmental toxicant exposures, the Toxicant Exposures and Responses by Genomic and Epigenomic Regulators of Transcription II (TaRGET II) Consortium was established in 2016^23^. This initiative systematically examined how early-life exposures to a broad range of environmental toxicants affect gene regulation across multiple tissues. The TaRGET II project aims to build a comprehensive reference atlas of environmentally induced molecular alterations, focusing on widely encountered toxicants such as arsenic (As), lead (Pb), endocrine-disrupting chemicals including bisphenol A (BPA), tributyltin (TBT), and di-2-ethylhexyl phthalate (DEHP), as well as dioxin (TCDD) and fine particulate matter (PM2.5). Maternal exposures were delivered via oral and inhalation routes, spanning from preconception through lactation, and high-throughput omics profiling of transcriptomic and epigenomic changes were then performed at three key life stages: weaning (∼3 weeks), early adulthood (5 months), and late adulthood (10 months) (Fig. 1A).

**Figure 1.**
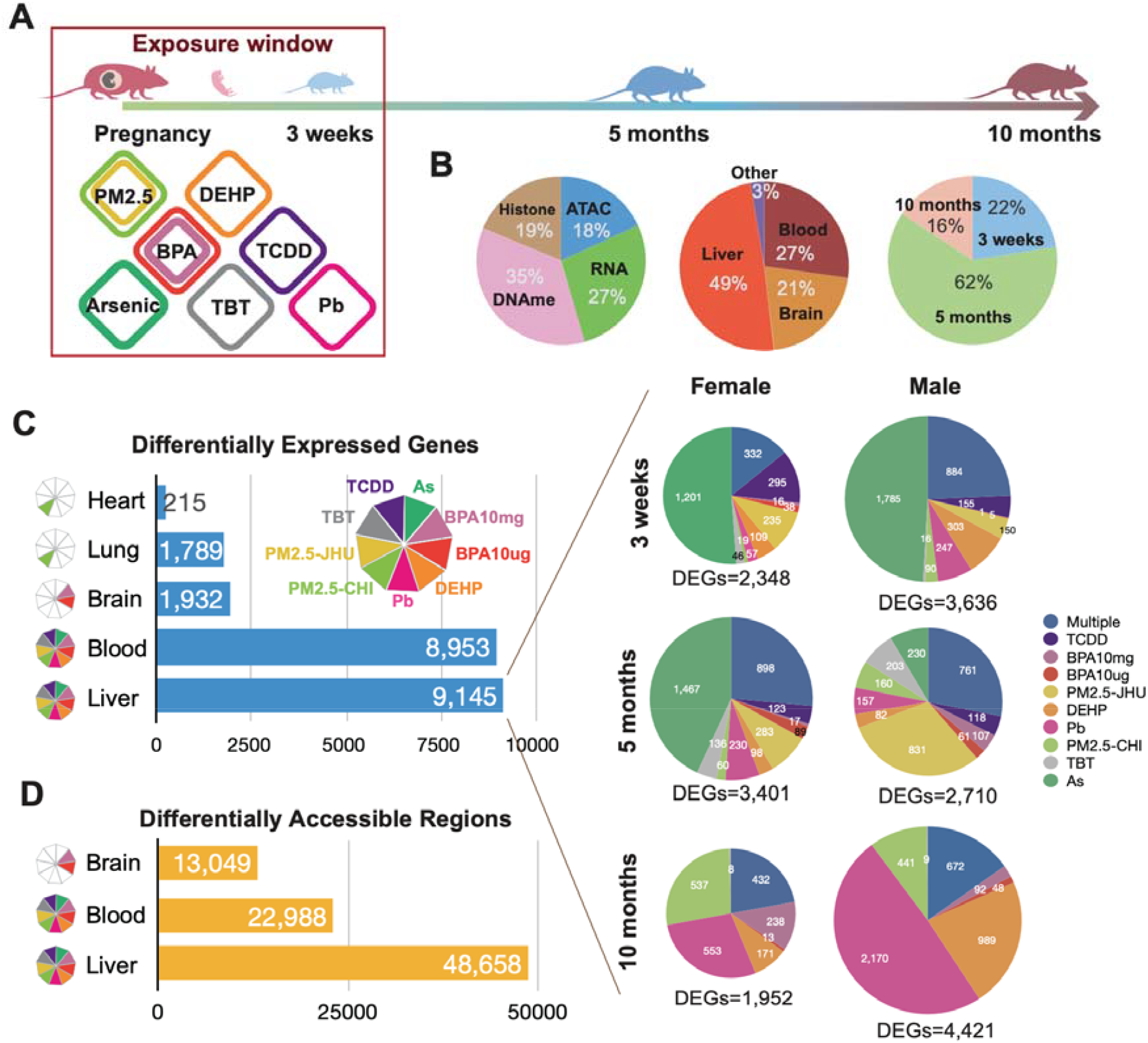
Summary of TaRGET II data production and exposure-specific molecular signatures. **A).** Experimental scheme for assessing the impact of early-life exposure on mouse development. **B)**. TaRGET II dataset statistics. **C)**. Left: Exposure-induced Differentially Expressed Genes (DEGs) in five tissues. Right: Exposure-specific DEGs in the mouse liver at three developmental stages. **D)**. Exposure-induced Differentially Accessible Regions (DARs) in three tissues.

In this article, we introduce an integrative multi-omics resource platform, ToxiTaRGET (https://toxitarget.com), which thoroughly combines and visualizes exposure-specific molecular signatures from the TaRGET II dataset. This includes both gene expression and epigenetic signatures across three different life stages (weaning, early, and late adulthood) in various tissues (blood, brain, liver, lungs, and ear) of both female and male mice. ToxiTaRGET allows researchers to query and explore toxicant-induced molecular changes, offering a valuable resource to understand how different tissues respond to various toxicant exposures by altering gene expression and remodeling the epigenetic landscape.

### ToxiTaRGET Data Statistics

The TaRGET II dataset contains over 3,000 epigenetic and transcriptomic profiles of distinct tissues at three developmental stages in mice (Fig. 3B). As the primary target tissue, the liver has comprehensive coverage across all early-life exposures, accounting for 49% of the total TaRGET II dataset. Using this dataset, we systematically identified differentially expressed genes (DEGs), differentially accessible regions (DARs), and differentially methylated regions (DMRs) for all tissues under various metadata conditions, including exposures, sexes, and developmental stages^23^. In the liver, 9,145 genes were identified as having altered expression in response to exposures. Across all developmental stages and sexes, exposure-induced DEGs consistently showed strong exposure-specific patterns (Fig. 1C). At 3 weeks, arsenic exposure induced the most significant gene expression changes in both sexes. In male livers, PM2.5-CHI exposure and lead (Pb) exposure induced the strongest gene expression changes at 5 months and 10 months old, respectively (Fig. 1C). We also identified distinct numbers of DARs and DMRs for different tissues (Fig. 1D, S-Table 1).

### ToxiTaRGET Database Access

ToxiTaRGET was created to host a comprehensive collection of molecular signatures that respond to various exposures in different tissues at three developmental stages in mice. The system provides quantitative measurements of genome-wide gene expression, ATAC-seq signals, and DNA methylation levels across all metadata conditions. These data are indexed using a MongoDB structure and linked to exposure-specific molecular signatures, including differentially expressed genes (DEGs), differentially accessible regions (DARs), and differentially methylated regions (DMRs). A PHP-based user interface was developed to enable real-time data queries, along with a JavaScript environment for visualizing gene expression data, ATAC-seq signals, and DNA methylation levels (Fig. 2A). ToxiTaRGET offers complete lists of exposure-specific molecular signatures identified from TaRGET II datasets, allowing users to browse and access these signatures directly [Fig.2B].

**Figure 2.**
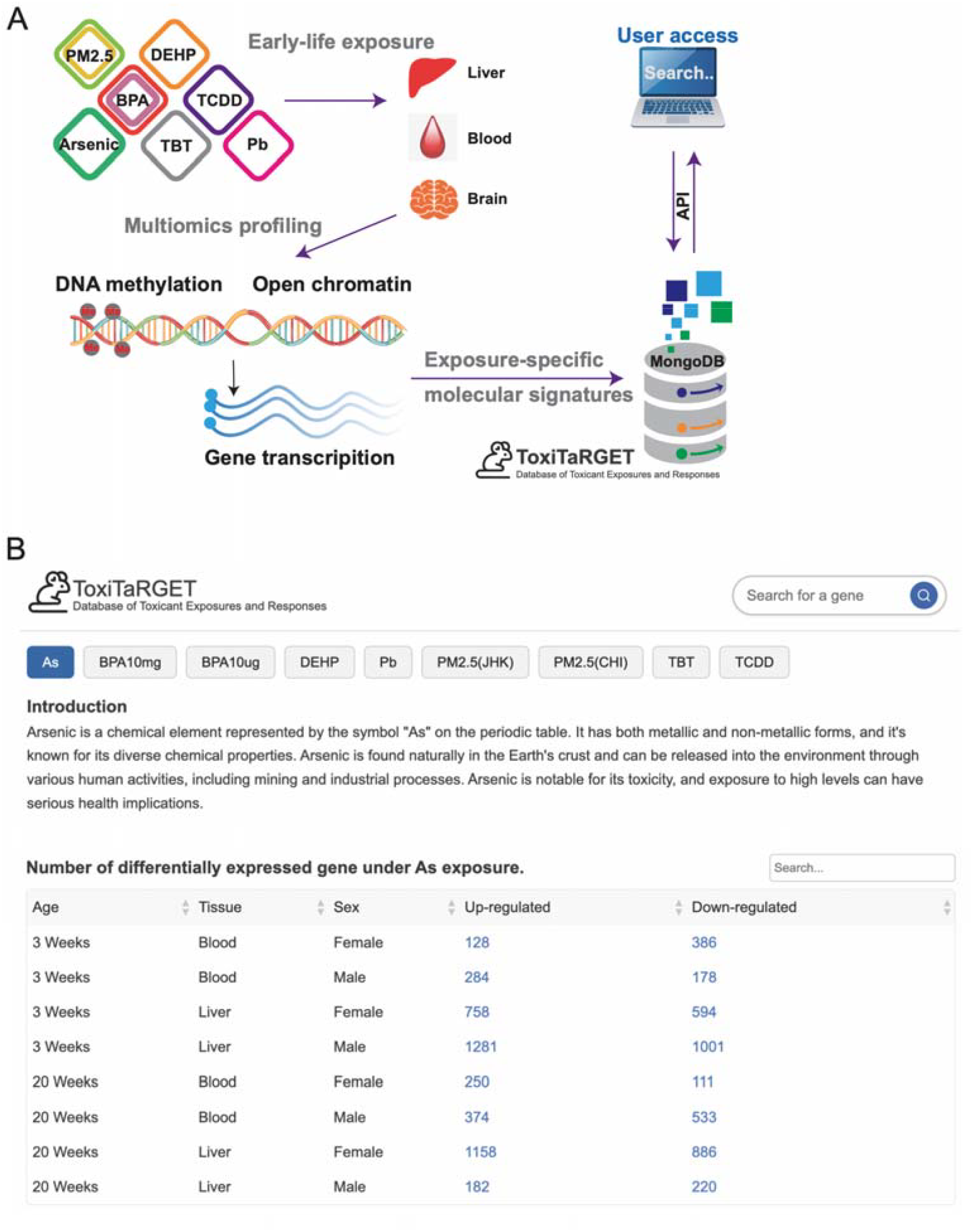
Identification and Collection of Multiomics Exposure-Specific Molecular Signatures. **A).** Scheme for Identification and Organization of Multiomics Exposure-Specific Molecular Signatures in the ToxiTaRGET Database. **B)**. Introduction and Summary of Arsenic-Specific DEGs Collection in Liver and Blood at 3 weeks and 5 months Stages

Meanwhile, ToxiTaRGET offers a user-friendly search interface, allowing users to easily query genes of interest by name (Fig. 3A). The search results are presented in a sectioned format, providing comprehensive outputs of gene expression, ATAC-seq signals, and DNA methylation levels all on a single page. For example, by searching for gene Gstm3, the “Gene Information” section includes the Ensembl ID, UniProt ID, gene status, structural visualization through AlphaFold, gene function, and subcellular location(Fig.3B). The gene function and subcellular location details were retrieved from UniProt^24^. The next section is derived from the RNA-Seq results and is divided into two sub-sections. The first sub-section indicates whether the queried gene is differentially expressed (DE) or non-differentially expressed across various exposures, tissues, and sexes (Fig.3C). This information is visually presented through an aesthetically pleasing heatmap, with each heatmap representing a specific sex. Both heatmaps incorporate data from all three age groups. The X-axis displays tissues and age groups, while the Y-axis illustrates exposures. Different colors indicate up- and down-regulated DEGs along with their fold changes. The second sub-section displays the expression pattern of the queried gene in various tissues under different exposures, relative to the control, using a box plot format (Fig.3C). For each sex, every tissue features two box plots encompassing all three age groups of mice. Different colors represent different ages. Users can choose to analyze a single age group or all ages simultaneously by utilizing the toggle button at the top right of each plot.

**Figure 3.**
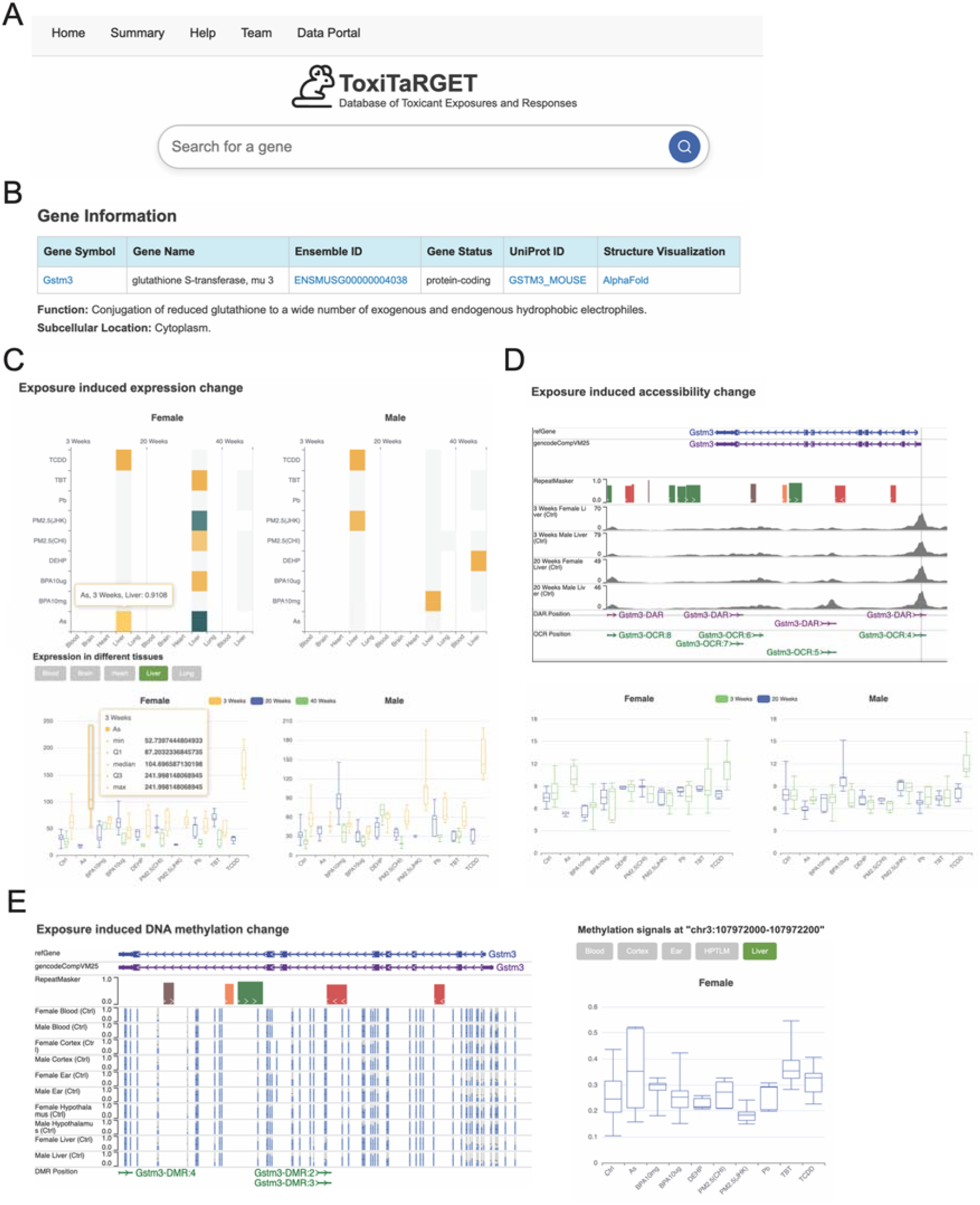
User Interface of ToxiTaRGET. A). Search Tool: Located at the top of the homepage. B). Gene Information Section: Displayed as part of the search results. C). Heatmap indicating differential expression of the searched gene across distinct exposures (y-axis) and tissues (x-axis). Box plots showing gene expression under different exposures (x-axis) at three developmental stages in the liver (Bottom). D). Genome browser view highlighting chromatin accessibility around the searched gene Gstm3 (Top). The bottom boxplot displays the ATAC-seq signal intensity of DARs located on the Gstm3 promoter under various exposures (x-axis). E). Genome browser view depicting DNA methylation around the searched gene Gstm3 (Left). Box plots displaying DNA methylation levels of DMRs around Gstm3 under different exposures (Right).

The following sections present chromatin accessibility data from ATAC-Seq results and DNA methylation analysis. Both epigenetic signatures are indexed based on their proximity to the nearest gene. In the ATAC-Seq results section, we integrated the WashU Epigenome Browser^25^, allowing users to access browser tracks for control conditions across all experimental scenarios, open chromatin regions (OCR), and differentially accessible regions (DAR) (Fig.3D). Clicking on specific OCR or DAR regions within the browser enables users to view the genomic region and the conditions under which these regions are identified as OCR or DAR. The second sub-section provides a table listing all OCRs and DARs associated with the queried gene. The OCRs are ordered by their distance from the transcription start site (TSS), from closest to farthest. Sorting options are available for each column, so users can organize the data according to their preferences. Each OCR in the table links to a dedicated page for detailed analysis, which includes two sub-sections: the fir^st^ features the WashU browser displaying control and exposure tracks for visual comparison; the second displays two box plots for each sex, showing the quantified ATAC-seq signal of OCRs in tissues of different ages under various exposures (Fig.3D). This box plot format closely resembles the one used in the RNA-Seq data section, offering users a familiar visualization style.

The final section shows results from DNA methylation data. A WashU browser displays control tracks for each tissue and sex, along with differentially methylated regions (DMRs) annotated with gene names (Fig.3E). Clicking on a DMR provides detailed information about its location on the chromosome, including exposure, tissue, age, sex, and hypo-or hyper-methylation status related to the queried gene. The second subsection is a table listing the DMRs associated with the queried gene. The default order is by decreasing proximity to the TSS. The table includes details about the DMR’s position, distance to TSS, tissue, sex, exposure, and methylation level, each in its own column. Sorting and search options are available for user convenience. Each DMR position is hyperlinked to a separate page with detailed insights, similar to the ATAC-seq dedicated page, including the DNA methylation plot for each DMR (Fig.3E).

## Discussion

The dynamic epigenome undergoes context-specific changes throughout the entire life span and various developmental stages^26-30^. Chronic exposure to environmental toxic chemicals can cause significant molecular changes, including shifts in gene transcription and epigenetic modifications. These alterations can lead to disease development and long-term health risks^31-33^ by reprogramming and disrupting specific gene expression programs that control biological functions in response to complex environmental stimuli^34-36^. With the rapid progress of toxicogenomic approaches, numerous high-throughput genomics datasets have been created to study molecular responses to toxicant exposures. These datasets provide valuable insights into toxicity mechanisms and disease risk. By the end of 2024, the NCBI Gene Expression Omnibus (GEO) contained 243,271 data series, representing 7.5 million biological samples. Such a vast dataset presents major challenges for researchers seeking to access specific data for particular biological questions.

To better understand how environmental toxicants impact gene regulation and cellular function, there is a growing need to integrate transcriptomic and epigenetic datasets. Over the past decade, several curated databases and repositories have emerged to support research in toxicogenomics, facilitating greater understanding of the molecular underpinnings of toxic exposures. The Comparative Toxicogenomics Database^37^ (CTD) is a literature-based curated repository that provides information on the relationships between chemicals, genes, and diseases. CTD also provided Tetramers^38^, a tool that explores CTD data and generates CGPD-tetramers, a four-unit block of information linking an initiating chemical, an interacting gene, a phenotype, and a disease outcome. While the CTD provides valuable information on gene expression changes under distinct toxicant exposures in various animals, it lacks detailed descriptions of gene expression. Additionally, CTD does not offer data on epigenetic modifications, nor does it provide insights into sex-specific or tissue-specific patterns. ToxicoDB^39^ is a cloud-based platform integrating data from large in vitro toxicogenomic studies, including gene expression profiles of primary human and rat hepatocytes treated with 231 potential toxicants. It provides users with harmonized chemical annotations, time- and dose-dependent plots of compounds across datasets, and toxicity-related pathway analysis, focusing on transcriptomic responses but not epigenetic changes. DrugMatrix^40^ database was one of the world’s largest toxicogenomic reference resources, and contains gene expression responses to environmental toxicant compound treatments in different tissues. However, the database is no longer updated, similar to TG-GATEs^41,42^ (Toxicogenomics Project-Genomics Assisted Toxicity Evaluation System), which provides toxicogenomic data from in vivo and in vitro studies, including transcriptomic data from liver and kidney tissues exposed to various compounds. T3DB^43^ is a resource with nearly 2900 common toxic substances along with detailed information on their chemical properties, descriptions, toxic effects, and gene targets with related expression datasets.

So far, ToxiTaRGET is the most distinctive platform for integrating and visualizing multi-omics data. It combines RNA-Seq, ATAC-Seq, and WGBS data from three different developmental stages of male and female mice after early-life exposure to various environmental toxicants: As, Pb, BPA, TBT, DEHP, TCDD, and PM2.5. Overall, ToxiTaRGET is a valuable tool that allows users to explore multi-omics data across different tissues, exposures, aging stages, and sexes in a customizable setup. ToxiTaRGET is a free, accessible resource that significantly advances the understanding of transcriptomic and epigenomic responses to environmental toxicant exposures in the scientific community.

## Material and methods

### Data collection and compilation

We used three sets of high-throughput sequencing data—RNA-Seq, ATAC-Seq, and WGBS—obtained from the TaRGET II Consortium^23^. The data captures changes in gene expression, chromatin landscape, and DNA methylation resulting from exposure to seven different environmental chemicals at three ages: 3 weeks, 20 weeks, and 40 weeks, in both male and female mice. It includes six key mouse tissues: Blood, Brain, Heart, Liver, Lung, Ear, and two brain sub-tissues (cortex and hypothalamus). For details, see https://target.wustl.edu/portal/.

### Raw sequence data and processing

RNA-seq data were processed as in previous studies^44,45^. Raw fastq files of RNA-seq data were processed by Cutadapt^46^ (v1.16; --quality-cutoff=15,10 --minimum-length=36), FastQC (v0.11.7), and STAR^47^ (v2.5.4b; --quantMode TranscriptomeSAM --outWigType bedGraph -- outWigNorm RPM) to do the trimming, QC report, and mouse genome mapping (mm10) using our own built pipeline, the TaRGET-II-RNA-seq-pipeline (https://github.com/Zhang-lab/TaRGET-II-RNAseq-pipeline). Then, gene expressions across normal and exposure samples were calculated by featureCounts^48^ (v1.5.1) within the pipeline based on GENCODE vM20 gene annotation of mouse genome.

Raw fastq of ATAC-seq data were processed by our own built pipeline based on AIAP^49^, the TaRGET-II-ATAC-seq-pipeline (https://github.com/Zhang-lab/TaRGET-II-ATACseq-pipeline) that integrated AIAP packages containing optimized QC reports and analysis pipeline with default parameters to generate the open chromatin regions (OCRs). Then, the consensus regions of OCRs across all exposure and control samples were generated with Index (https://github.com/Altius/Index) method used for the downstream analysis. The ATAC-seq signals of consensus OCRs were calculated by using the intersectBed method.

The TaRGET-II-WGBS-pipeline (https://github.com/Zhang-lab/WGBS_analysis) was built by using the Cutadapt^46^ (v1.16; --quality-cutoff=15,10 --minimum-length=36) and Bismark^50^ (v0.19.0; --bowtie2 -X 1000 --score_min L,0,-0.6 -N 0 --multicore 2 -p 4) to do the trimming and mouse genome mapping. The pipeline also incorporated quality control, generated user-friendly files for computational analysis, and output genome browser tracks for data visualization.

### Normalization and Batch Effect Correction for RNA-seq and ATAC-seq Data

RNA-seq and ATAC-seq datasets were normalized to correct for batch effects arising from different data production centers. In general, RNA-seq and ATAC-seq data generated within the same production center were analyzed together. Gene counts from RNA-seq were first normalized using the relative log expression (RLE) method implemented in *edgeR*. Subsequently, factor analysis was performed across exposure groups by applying the residuals calculation of the *RUVr* function in *RUVSeq*^*51*^. To further remove unwanted technical variations (k=3) unrelated to exposures, including biases from tissue dissection and library preparation, a general linear model fitting process was employed as in previous studies^44,45^.

### Differentially analysis

Differentially expressed genes (DEGs) between control and exposure samples under varying conditions were identified using DESeq2^52^ (v1.34.0) as previous studies^53-55^. Consortium-normalized read count data were used to identify the DEGs between the exposed and control samples. Genes were considered significantly differentially expressed if they met the following stringent criteria: an adjusted p-value (Benjamini-Hochberg correction) less than 0.001 and an absolute log2 fold change greater than log2(1.5), equivalent to a 1.5-fold change in expression.

Differentially accessible regions (DARs) between control and exposure samples were identified using edgeR^56^ (v3.36.0), as previous study^49,57,58^. Consortium-normalized ATAC-seq reads were used to identify DARs, with significance determined by the following cutoffs: an absolute log2 fold change greater than log2(1.5) (a 1.5-fold change in accessibility) and a false discovery rate (FDR) less than 0.01. These criteria balance sensitivity and specificity, ensuring robust detection of chromatin accessibility changes associated with exposure.

Differential methylation regions (DMRs) between control and exposure samples were identified using a method adapted from previous studies^27,28,59,60^. Methylation levels were quantified using whole-genome bisulfite sequencing (WGBS) data. Briefly, raw WGBS reads were aligned to the reference genome using Bismark^50^ (v0.23.1) with default parameters, followed by deduplication to remove PCR artifacts. Methylation levels at individual CpG sites were calculated as the ratio of methylated reads to total reads (methylated + unmethylated) at each site, with a minimum coverage of 10 reads per site to ensure reliability. DMRs were identified through a genome sliding approach, as previous studies^59-61^: the genome was divided into sliding 200bp windows, the methylated and unmethylated reads counts in each window were calculated in both control samples and exposed samples, and then a Chi-squared test was applied to identify DMRs. Regions were defined as differentially methylated if they exhibited a methylation level difference of at least 0.1 (a 10% absolute difference in methylation between control and exposed condition) and a q-value (adjusted for multiple testing using the BH method) less than 0.1. Additionally, only regions with at least 2 CpG sites and a minimum average coverage of 20 reads per site were considered to enhance statistical power and reduce noise in the analysis, and the neighboring regions were merged to maximize the width of DMRs.

### Web interface and database architecture

The backend of the platform was built on a Google Cloud-based Apache server (Apache 2.4.58 ; Ubuntu 18.04 LTS) using the Flask framework (3.1.0; https://palletsprojects.com/), which facilitated API routing, data processing, and communication between the front-end and the database. MongoDB (https://www.mongodb.com/) was utilized to manage the retrieval and storage of structured and semi-structured data, including gene expression profiles and associated metadata. Flask handled requests from the front-end, processed data, and interfaced with MongoDB for efficient data management through PyMongo (4.11.3; https://pymongo.readthedocs.io/).

On the front-end, the React framework (v19.1; https://react.dev/) was employed to develop the user interface, delivering a responsive and interactive experience for users. The JavaScript library Apache ECharts (5.6; https://echarts.apache.org/) was integrated to support dynamic visualization, allowing for the comprehensive and real-time display of complex data sets. The database schema and the relationships among its components, focusing on gene expression profiles, metadata, and other structured information, are illustrated in Figure 1. This schematic provides an overview of how data flows through the system, from storage in MongoDB to visualization in the user interface.

## DATA Availability

https://target.wustl.edu/portal/.

## Acknowledgements

This work was supported by the National Institutes of Health: R35GM142917, U24ES026699, U24HG012070, U24NS132103, U01ES026719, P30ES030285. Funding for the open-access charge was provided by the National Institutes of Health.

## Supplementary data

Conflict of interest statement. None declared.

